# Inhibition of Human Mantle Cell Lymphomas by the β_2_-Adrenergic Agonist, Levalbuterol, in the Hollow Fiber Assay in Mice

**DOI:** 10.1101/2025.09.01.673249

**Authors:** Mario A. Inchiosa

## Abstract

This study follows from the novel findings in our previous publication that β_2-_adrenergic receptor (β2AR) agonists show gene-expression signatures that are consistent with glucocorticoid receptor (GR) agonists. These observations were made with analytical software associated with the Harvard/MIT genomic database, CLUE. This formed a hypothesis that β2AR agonists would share the anti-inflammatory activity of GR agonists and their ability to induce apoptosis in tumors of lymphatic lineage. Our lead β2AR agonist from our previous study, levalbuterol, was tested for inhibitory activity upon 3 non-Hodgkin human cell lines, two of which were Mantle Cell carcinomas, JEKO-1 and MINO, and against SU-DHL-1, in mice: Species: *Mus musculus;* Strain: NMRI nude (Crl:NMRI-Foxn1nu), Sex: Females. The drug was supplied ad libitum in the drinking water at concentrations of 8 µg/ml and16 µg/ml over a period of 14 days. Statistically significant inhibition of lymphoma proliferation was observed at the higher concentration of levalbuterol. The data for the 3 cultures were combined to increase the power of the analysis. The average inhibition was 55 %, p <0.0001, consistent with the hypothesis that the β2AR agonist shared the apoptotic activity of the GR agonists with which it had similar gene-expression activity.

## 1. Introduction

The present study has its foundation from the novel observation that β_2_-adrenergic receptor (β2AR) agonists have gene-expression signatures that are remarkably similar to those of glucocorticoid receptor (GR) agonists [1]. These observations were made through analyses with the Harvard/MIT Broad Institute genomic database, CLUE. When the prototypical β2AR agonist epinephrine was profiled on CLUE it was found to have scores for gene-expression that were highly similar to those of GR agonists.

Some of the evidence for this similarity between gene-expression of β2AR agonists and GR agonists is presented here for perspective.

Table 5 from reference No. 1 shows the gene-expression connectivity (“similarity”) scores for epinephrine vs GR agonists.

**Table 5.** Gene-Expression Connectivity Scores for EPINEPHRINE vs GR Agonists.^*^

|  |  | Rank | ScoreName | Description |
| --- | --- | --- | --- | --- |
| 4 | 99.89 |  | hydrocortisone | Glucocorticoid receptor agonist |
| 5 | 99.89 |  | betamethasone | Glucocorticoid receptor agonist |
| 7 | 99.79 |  | budesonide | Glucocorticoid receptor agonist |
| 8 | 99.79 |  | flunisolide | Cytochrome P450 inhibitor |
| 9 | 99.75 |  | mometasone | Glucocorticoid receptor agonist |
| 11 | 99.75 |  | fluocinonide | Glucocorticoid receptor agonist |
| 13 | 99.68 |  | triamcinolone | Glucocorticoid receptor agonist |
| 14 | 99.68 |  | fludroxycortide | Glucocorticoid receptor agonist |
| 19 | 99.62 |  | clobetasol | Glucocorticoid receptor agonist |
| 21 | 99.61 |  | triamcinolone | Glucocorticoid receptor agonist |
| 22 | 99.61 |  | flumetasone | Glucocorticoid receptor agonist |
| 23 | 99.58 |  | fluticasone | Glucocorticoid receptor agonist |
| 24 | 99.58 |  | fluorometholone | Glucocorticoid receptor agonist |
| 26 | 99.54 |  | dexamethasone | Glucocorticoid receptor agonist |
| 27 | 99.52 |  | prednicarbate | Phospholipase activator |
| 28 | 99.51 |  | isoflupredone | Glucocorticoid receptor agonist |
| 29 | 99.47 |  | hydrocortisone | Glucocorticoid receptor agonist |
| 30 | 99.44 |  | diflorasone | Corticosteroid agonist |
| 31 | 99.44 |  | dexamethasone | Glucocorticoid receptor agonist |
| 32 | 99.4 |  | medrysone | Glucocorticoid receptor agonist |
| 33 | 99.37 |  | prednisolone | Glucocorticoid receptor agonist |
| 34 | 99.33 |  | halcinonide | Glucocorticoid receptor agonist |
| 37 | 99.12 |  | betamethasone | Glucocorticoid receptor agonist |
| 39 | 99.05 |  | beclometasone | Glucocorticoid receptor agonist |
| 40 | 99.05 |  | fluocinolone | Glucocorticoid receptor agonist |
| 42 | 99.05 |  | hydrocortisone | Glucocorticoid receptor agonist |
| 43 | 99.01 |  | hydrocortisone | Glucocorticoid receptor agonist |
| 45 | 98.77 |  | clocortolone | Glucocorticoid receptor agonist |
| 46 | 98.77 |  | beclometasone | Glucocorticoid receptor agonist |
| 48 | 98.6 |  | westcort | Glucocorticoid receptor agonist |
| 51 | 98.4 |  | prednisolone | Glucocorticoid receptor agonist |
| 52 | 98.38 |  | fluticasone | Glucocorticoid receptor agonist |
| 55 | 98.34 |  | hydrocortisone | Glucocorticoid receptor agonist |
| 56 | 98.32 |  | rimexolone | Glucocorticoid receptor agonist |
| 58 | 98.17 |  | fludrocortisone | Glucocorticoid receptor agonist |
| 59 | 98.17 |  | amcinonide | Glucocorticoid receptor agonist |
| 61 | 98.03 |  | methylprednisolone | Glucocorticoid receptor agonist |
| 67 | 97.71 |  | desoximetasone | Glucocorticoid receptor agonist |
| 75 | 97.67 |  | halometasone | Glucocorticoid receptor agonist |
| 88 | 97.53 |  | loteprednol | Glucocorticoid receptor agonist |
| 96 | 97.5 |  | depomedrol | Glucocorticoid receptor agonist |
| 117 | 97.39 |  | prednisolone | Glucocorticoid receptor agonist |
| 137 | 97.15 |  | fluocinonide | Glucocorticoid receptor agonist |
| 632 | 93.24 |  | alclometasone | Glucocorticoid receptor agonist |

\*From reference No. 1

It should be noted that gene-expression connectivity scores above 90 are considered to have potential biological importance that may be worth consideration for further investigation.

Quite remarkably, Table 9 below from reference No. 1 shows the gene-expression connectivity scores for cortisol (hydrocortisone) vs GR agonists; they profile in scoring almost identically to those for epinephrine.

**Table 9.** Gene-expression connectivity scores for HYDROCORTISONE vs GR agonists.^*^

|  | Rank | Score | Name | Description |
| --- | --- | --- | --- | --- |
| 1 | 99.99 |  | hydrocortisone | Glucocorticoid receptor agonist |
| 2 | 99.93 |  | triamcinolone | Glucocorticoid receptor agonist |
| 3 | 99.93 |  | budesonide | Glucocorticoid receptor agonist |
| 4 | 99.93 |  | fluocinonide | Glucocorticoid receptor agonist |
| 5 | 99.93 |  | fluorometholone | Glucocorticoid receptor agonist |
| 7 | 99.72 |  | mometasone | Glucocorticoid receptor agonist |
| 8 | 99.72 |  | fluticasone | Glucocorticoid receptor agonist |
| 9 | 99.72 |  | fludrocortisone | Glucocorticoid receptor agonist |
| 11 | 99.61 |  | betamethasone | Glucocorticoid receptor agonist |
| 12 | 99.61 |  | betamethasone | Glucocorticoid receptor agonist |
| 13 | 99.58 |  | clocortolone | Glucocorticoid receptor agonist |
| 14 | 99.58 |  | flunisolide | Cytochrome P450 inhibitor |
| 15 | 99.58 |  | dexamethasone | Glucocorticoid receptor agonist |
| 16 | 99.58 |  | clobetasol | Glucocorticoid receptor agonist |
| 17 | 99.54 |  | isoflupredone | Glucocorticoid receptor agonist |
| 20 | 99.37 |  | fludroxycortide | Glucocorticoid receptor agonist |
| 21 | 99.33 |  | prednisolone | Glucocorticoid receptor agonist |
| 22 | 99.3 |  | beclo methasone | Glucocorticoid receptor agonist |
| 23 | 99.3 |  | fluticasone | Glucocorticoid receptor agonist |
| 24 | 99.3 |  | alclometasone | Glucocorticoid receptor agonist |
| 26 | 99.3 |  | hydrocortisone | Glucocorticoid receptor agonist |
| 27 | 99.26 |  | fluocinolone | Glucocorticoid receptor agonist |
| 28 | 99.25 |  | flumetasone | Glucocorticoid receptor agonist |
| 30 | 99.16 |  | rimexolone | Glucocorticoid receptor agonist |
| 32 | 99.06 |  | loteprednol | Glucocorticoid receptor agonist |
| 34 | 98.91 |  | hydrocortisone | Glucocorticoid receptor agonist |
| 35 | 98.8 |  | halcinonide | Glucocorticoid receptor agonist |
| 36 | 98.66 |  | triamcinolone | Glucocorticoid receptor agonist |
| 37 | 98.63 |  | dexamethasone | Glucocorticoid receptor agonist |
| 38 | 98.59 |  | beclo methasone | Glucocorticoid receptor agonist |
| 39 | 98.52 |  | prednicarbate | Phospholipase activator |
| 44 | 98.48 |  | prednisolone | Glucocorticoid receptor agonist |
| 45 | 98.48 |  | amcinonide | Glucocorticoid receptor agonist |
| 46 | 98.1 |  | halometasone | Glucocorticoid receptor agonist |
| 48 | 97.82 |  | methylprednisolone | Glucocorticoid receptor agonist |
| 50 | 97.68 |  | diflorasone | Corticosteroid agonist |
| 52 | 97.5 |  | hydrocortisone | Glucocorticoid receptor agonist |
| 53 | 97.5 |  | fluocinonide | Glucocorticoid receptor agonist |
| 55 | 97.36 |  | desoximetasone | Glucocorticoid receptor agonist |
| 58 | 97.33 |  | prednisolone | Glucocorticoid receptor agonist |
| 66 | 97.15 |  | depomedrol | Glucocorticoid receptor agonist |
| 83 | 96.69 |  | medrysone | Glucocorticoid receptor agonist |
| 141 | 95.45 |  | westcort | Glucocorticoid receptor agonist |
| 187 | 95.1 |  | hydrocortisone | Glucocorticoid receptor agonist |

\*From reference No. 1

The association between the data for epinephrine and hydrocortisone vs GR agonists is emphasized by the presentation in Figure 4 from reference No. 1, below.

Several agents that are related to beta2-adrenergic agonists showed GR agonist activity when profiled on CLUE. However, the profound uniqueness of the strength of the gene-expression association between epinephrine and hydrocortisone as GR agonists demonstrate this association when all members of the group are compared as in Figure 5 in reference No. 1.

On the basis of the similarity of gene-expression for epinephrine and hydrocortisone vs GR agonists, it was hypothesized that the β2AR agonist, levalbuterol, may have the apoptotic activity of GR agonists and inhibit proliferation of malignancies of lymphatic lineage. This concept was tested in the present study against 3 non-Hodgkin human cell lines, two of which were Mantle Cell carcinomas, JEKO-1 and MINO, and against SUDHL-1, in mice.

### 1.1. Notes on the Hollow Fiber Assay Model

The cell cultures were placed in sections of straw-like hollow fibers of polyvinylidene fluoride of one millimeter diameter with 500kDa cut-off pores that restrict passage of cells but allow drugs, nutrients, oxygen, etc. to pass freely. Three, 2- cm sections were placed both intraperitoneally (i.p.) and subcutaneously (s.c.), in the neck region, according to the hollow fiber model developed by the United States National Cancer Institute for anti-tumor drug screening. Each animal received the 3 tumor cultures in individual sections both i.p. and s.c.. The model has been extensively studied and applied [2-6].

Levalbuterol was supplied ad libitum in the drinking water at concentrations of 0 (vehicle controls) and 8 and 16 µg/ml to 3 groups of 6 mice for 14 days. The number of live mice remaining was determined by measurement of ATP content (see **Materials and Methods**). Inhibition of tumor cell proliferation was only seen at the higher drug concentration. The data for the 3 lymphoma cultures were combined to increase the statistical power of the analyses; the average inhibition was 54.77 ± 5.69 (SEM) percent; p <0.0001.

## Results

### 2.1. Animal Body Weight Changes; Tumor Proliferation Rates and Inhibition by Levalbuterol

Average body weight changes in the vehicle control and 16 µg/ml levalbuterol treatment group during the experimental procedure are recorded in Figure 1.

**Figure 1.**
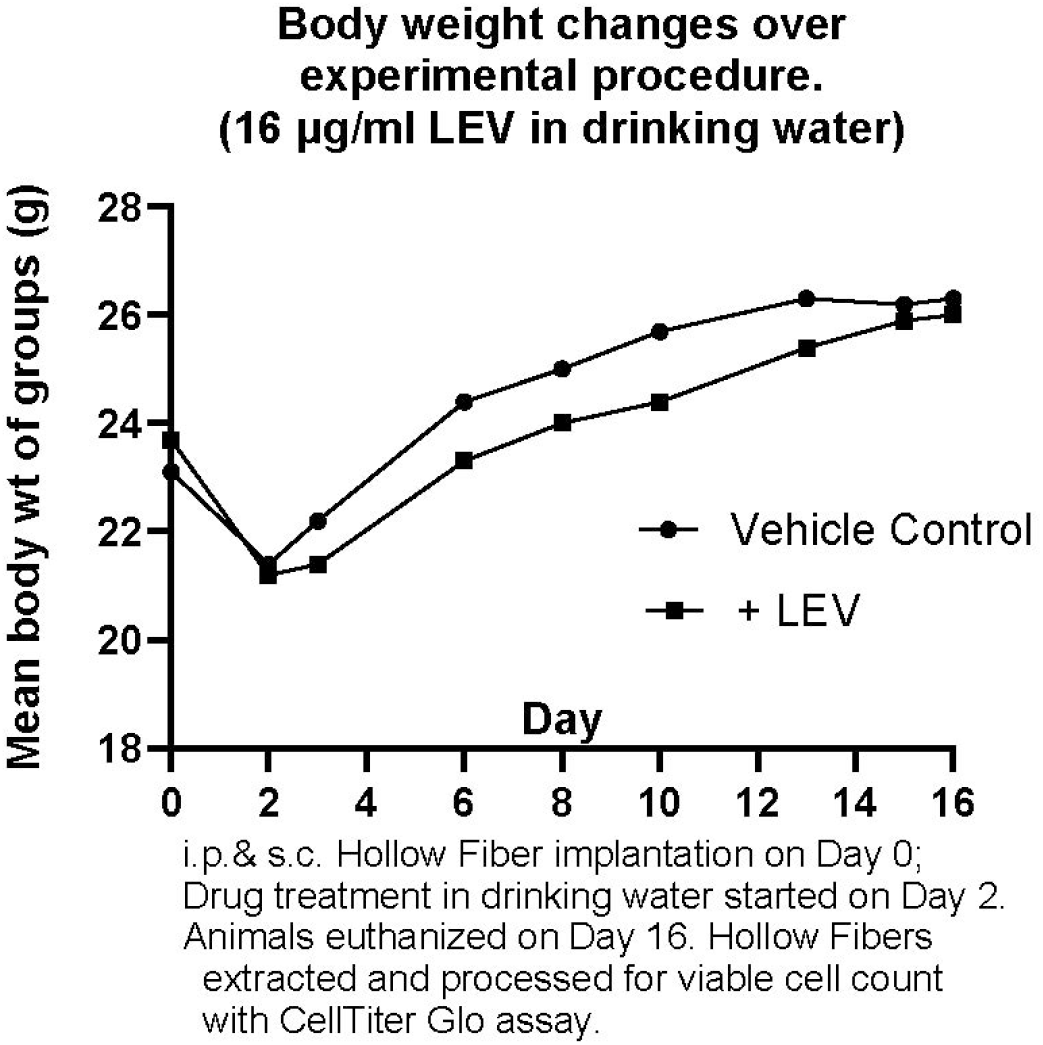
Body weght changes of groups at 16 µg/ml levalbuterol.

It appears that animal weight was well maintained over the experimental period.

Figure 2 presents the descriptive data for tumor cell proliferation for a levalbuterol drug treatment of 8 µg/ml over the 14-day study period.

**Figure 2.**
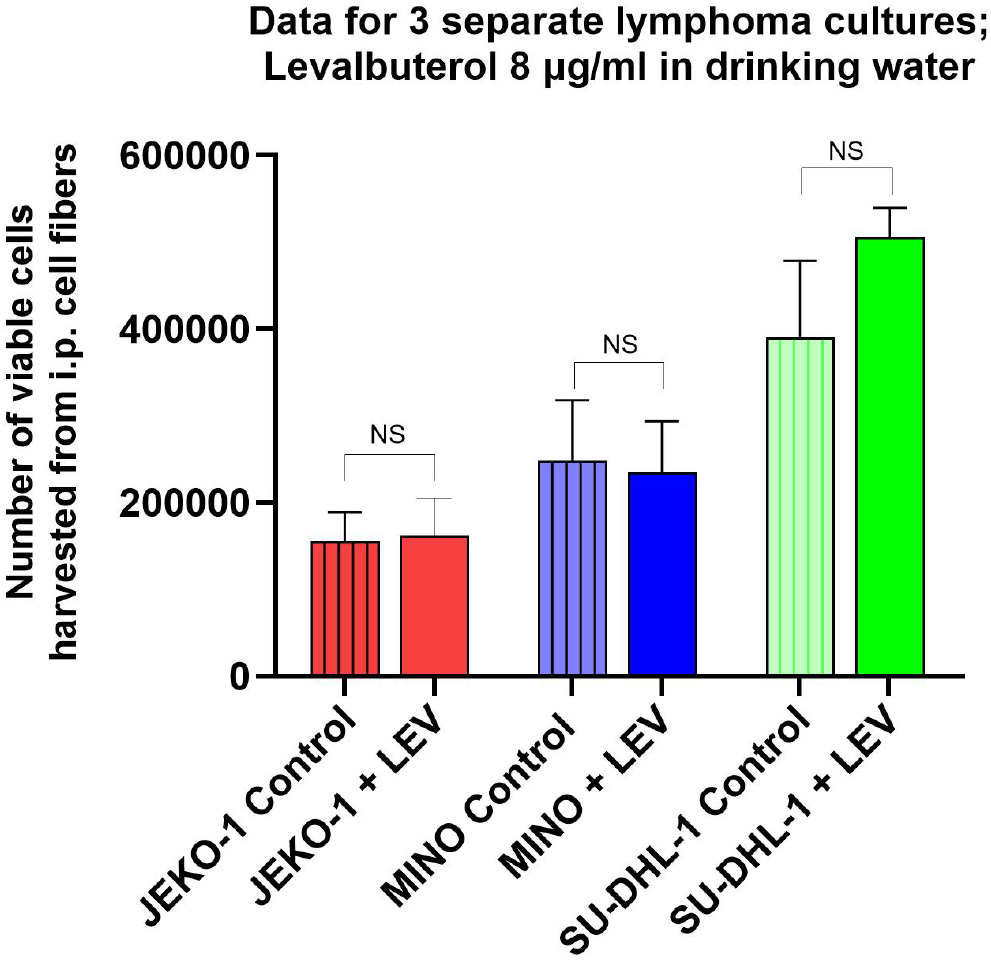
A treatment concentration of levalbuterol of 8 µg/ml in the drinking water did not result in any effect on proliferation in the malignant cell cultures. The data show the differences in proliferation rates among the 3 lymphoma cultures. The data show mean values and SEM.

Figure 3 presents the descriptive data for tumor cell proliferation for a levalbuterol drug treatment of 16 µg/ml over the 14 day study period.

**Figure 3.**
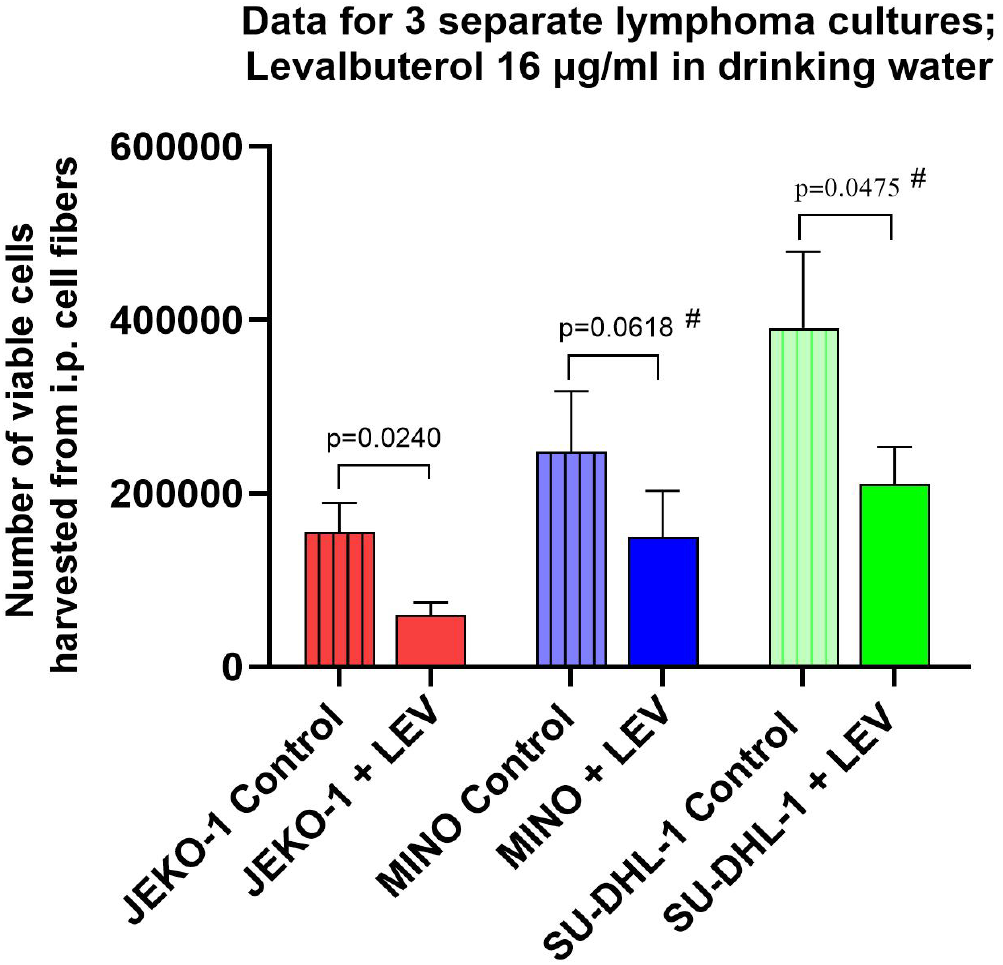
A treatment concentration of 16 µg/ml of levalbuterol in the drinking water resulted in inhibition of proliferation of all 3 non-Hodgkin lymphoma cultures to variant degrees. JEKO-1 was statistically inhibited in a two-sided unpaired t test. MINO trended closely to significance in a onesided unpaired t test (#) and SU-DHL-1 was significant in a one-sided unpaired t test (#). The data show mean values and SEM.

To increase the statistical power of the inhibitory effect of levalbuterol on the proliferation of the non-Hodgkin tumor cells, the data for the 3 cultures were combined. In order to accomplish this, it was necessary to use statistical methods to normalize the data to account for the intrinsic differences among the proliferation rates of the three tumor cultures as seen in Figure 3. In the first test of this, a one sample t test was conducted. A percent change in viable cell number from that of its control mean value was calculated for each culture. The results of those calculations are shown in Figure 4 for the cultures incubated i.p. at a treatment concentration of 16 µg/ml.

**Figure 4.**
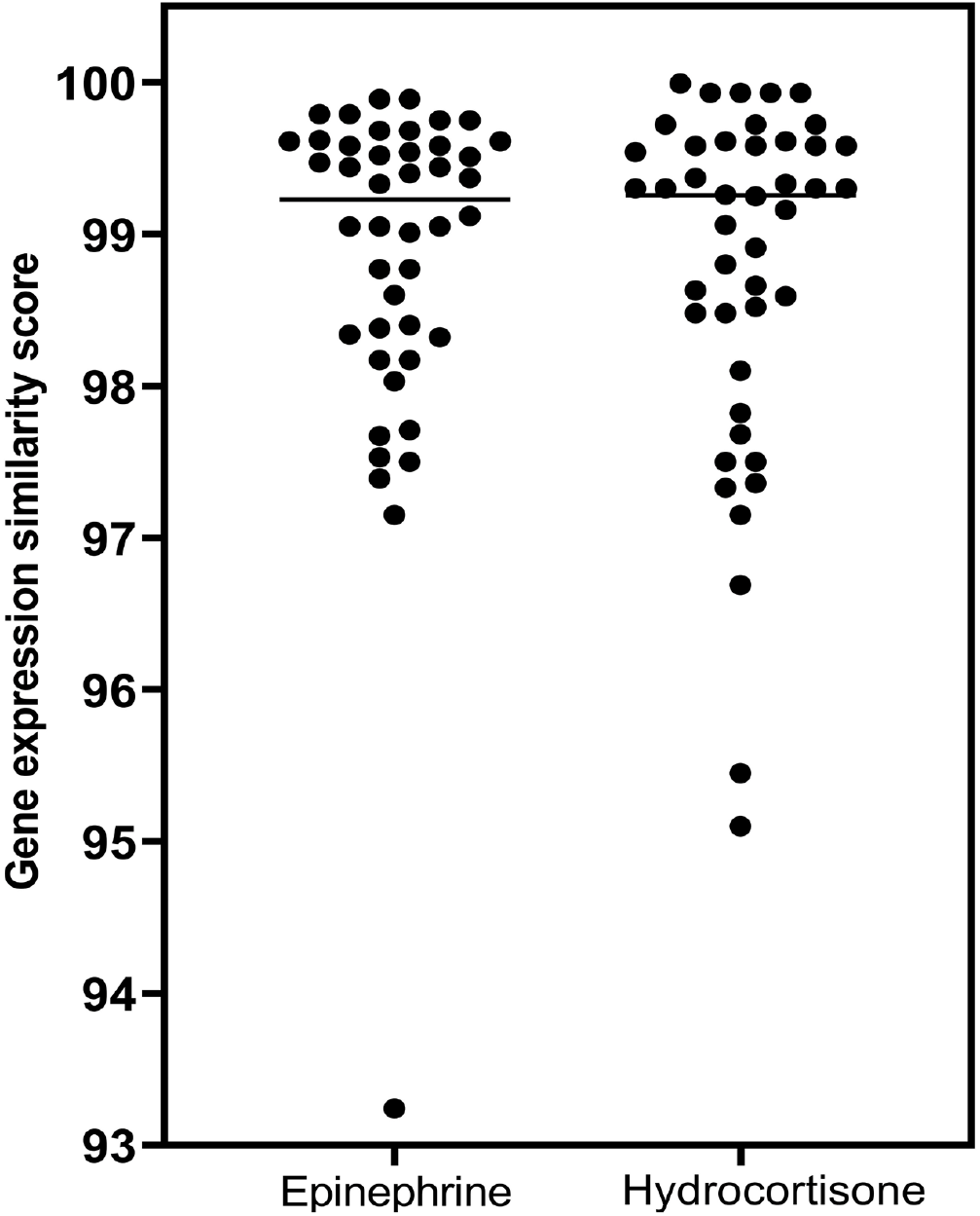
(From reference No. 1) Comparison of epinephrine and hydrocortisone gene-expression scores as glucocorticoid receptor agonists. The horizontal lines are the median values for the individual distributions, which are essentially identical. Mann-Whitney analysis of the distributions showed no difference in gene-expression characteristics for epinephrine and hydrocortisone as glucocorticoid receptor agonists (p = 0.805).

**Figure 4.**
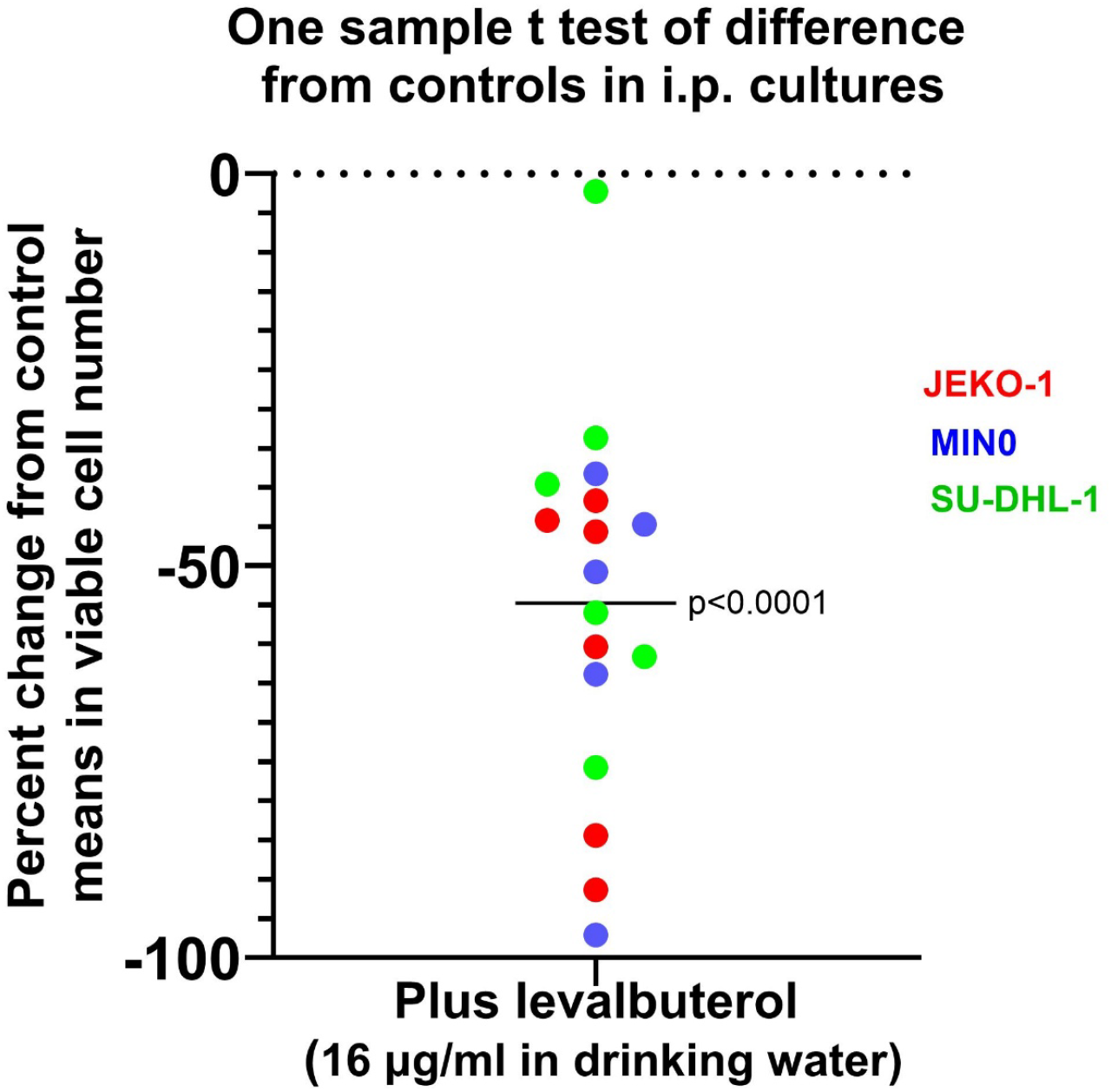
The overall average inhibition for the three cultures was 54.77 ± 5.69 (SEM) percent; p <0.0001.

A second statistical test was applied to normalize the same data for the inhibitory effect of levalbuterol on the i.p. non-Hodgkin lymphoma cultures. Again, this was necessary because the cultures had different intrinsic proliferation rates. In this case, the viable number of active cells was calculated as a fraction of its respective control mean for both the controls and for the cells that received levalbuterol at 16 µg/ml in the drinking water. This test represented a conventional unpaired t test; the results are presented in Figure 5.

**Figure 5.**
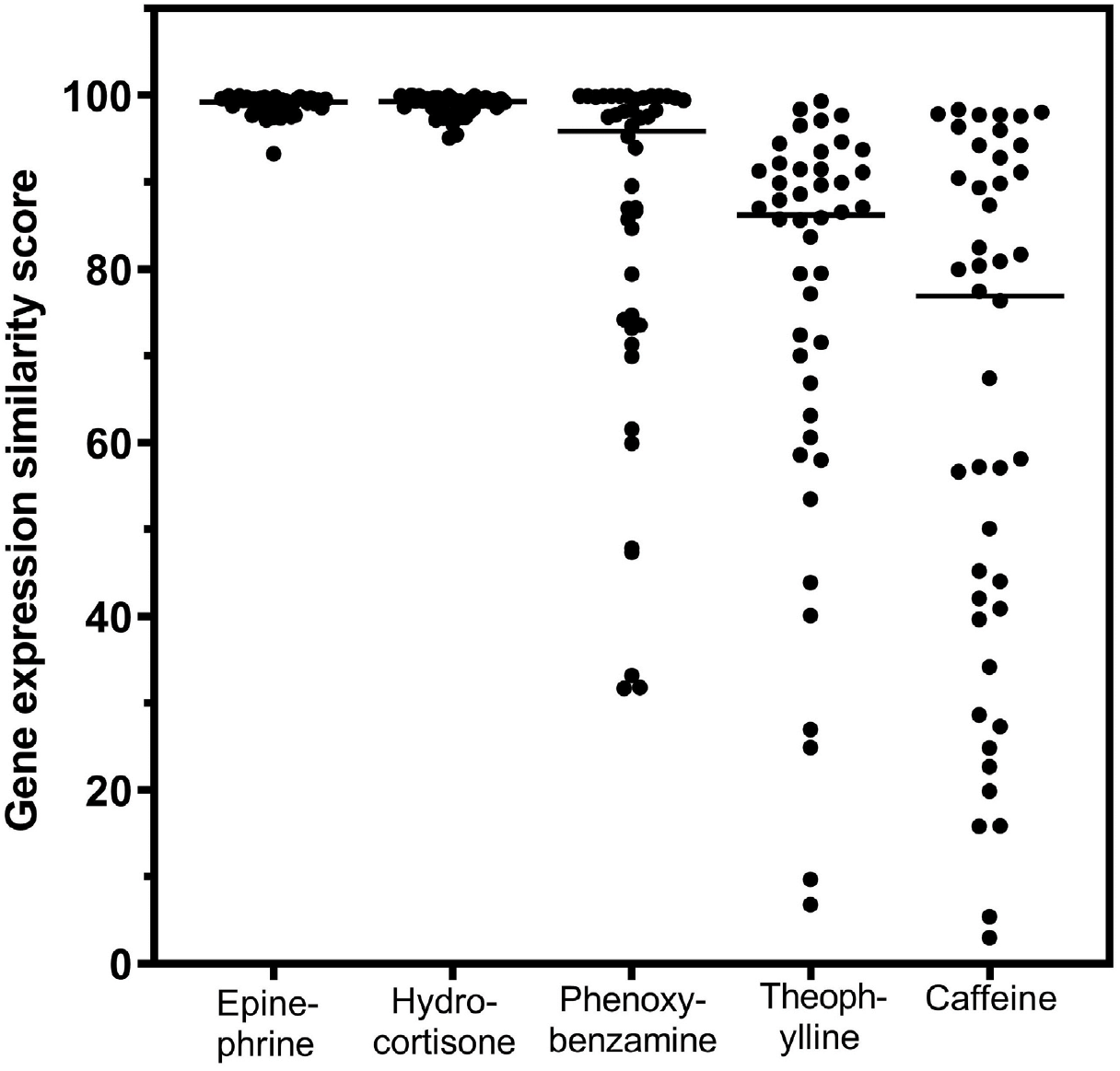
(From reference No. 1) Comparison of beta2-adrenergic related agents and hydrocortisone for gene expression connectivity scores as glucocorticoid receptor agonists. The horizontal lines are the median values for the individual distributions. The relative differences among the gene-expression scores can be appreciated visually. (Caffeine was not studied further in the investigations of reference 1.)

**Figure 5.**
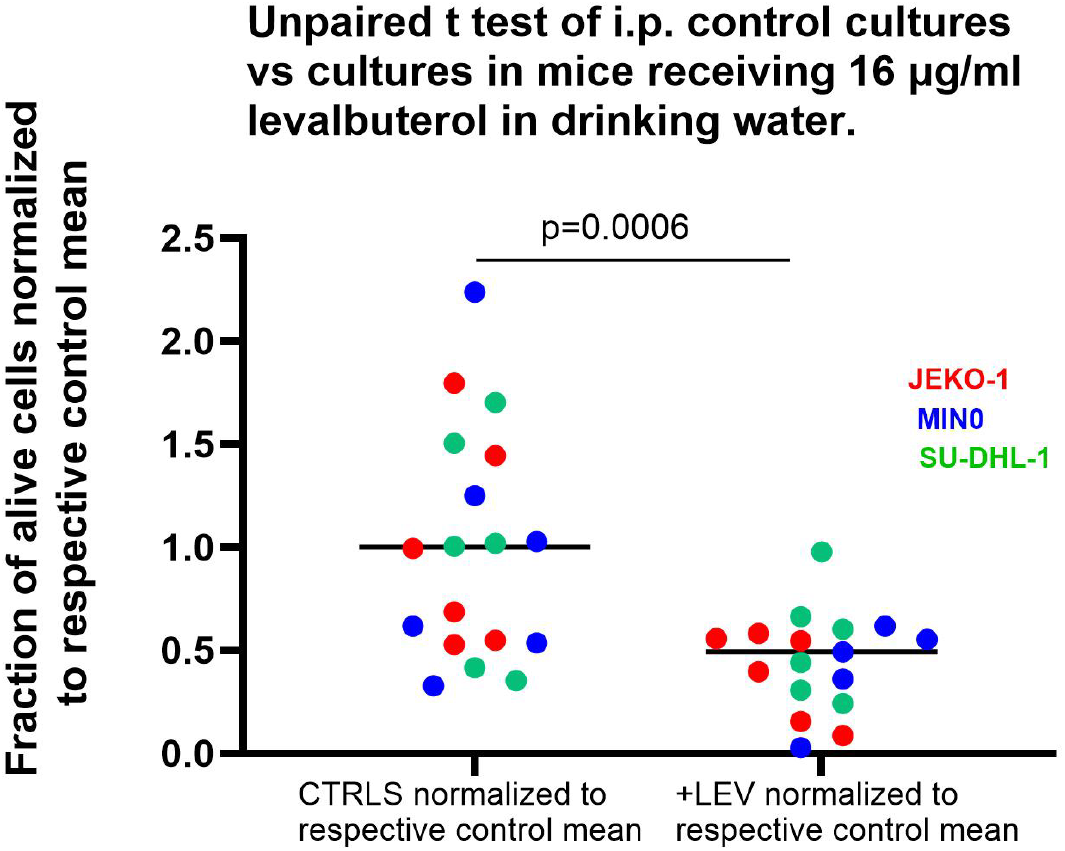
The overall average inhibition for the three cultures was 55.21 ± 14.63 (SEM) percent; p=0.0006. This is almost identical to the inhibition observed in the one sample t test presented for the same data in Figure 4. Both approaches to normalize the data were found to adjust for the different rates of proliferation among the 3 tumor cell cultures.

A one sample t test was conducted to analyze the inhibitory effect of levalbuterol in the s.c. cultures (Figure 6).

**Figure 6.**
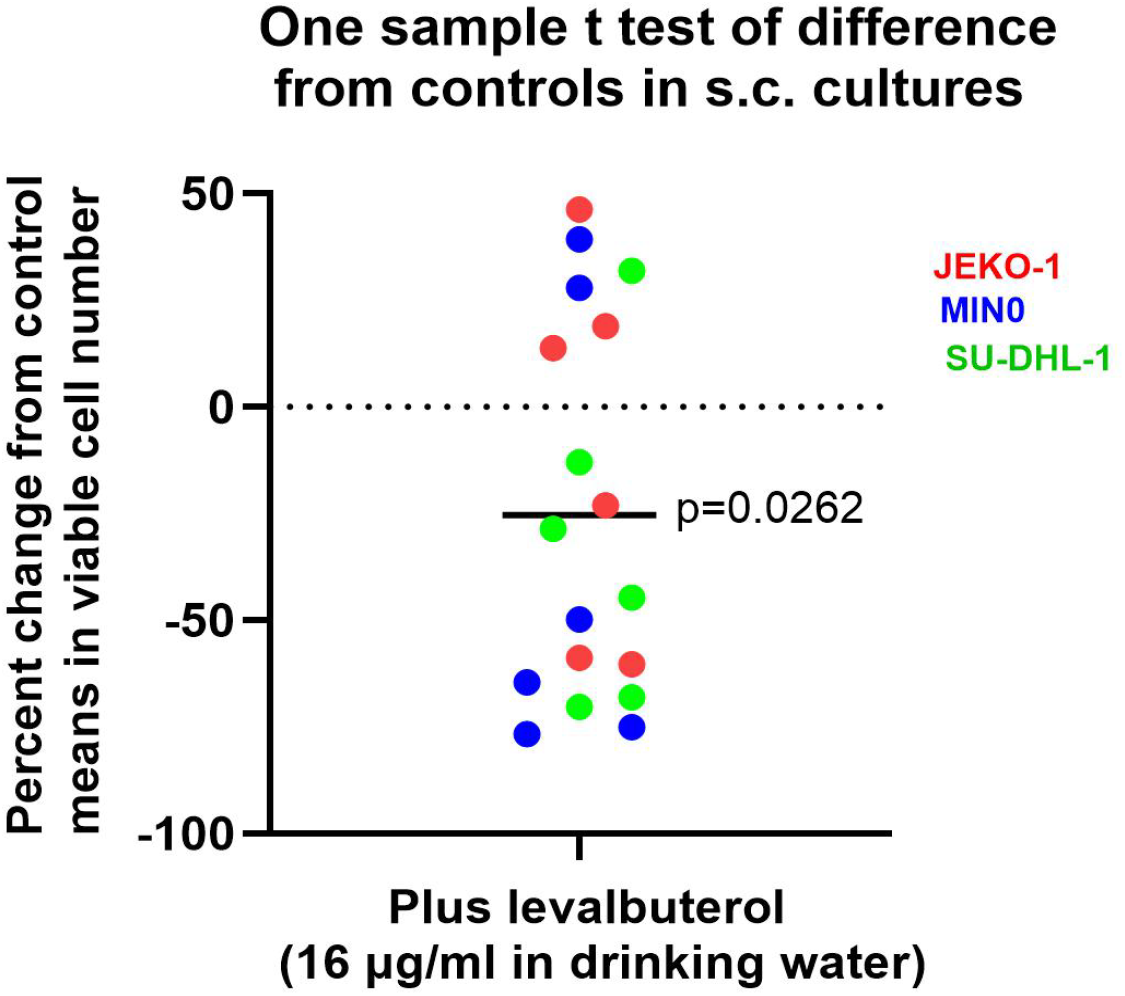
The overall average inhibition for the 3 cultures was 25.26 ± 10.37 (SEM) percent; p=0.0262. The inhibition from the s.c, compartment was not as strong as for i.p.; the access of levalbuterol to plasma bathing the s.c. compartment may have been less robust than i.p.

### 2.2. Toxicity Observations

A necropsy report was prepared by the contract research organization that conducted the Hollow Fiber assay (Reaction Biology Europe GmbH, Engesserstr.4, 79108 Freiburg, Germany). Their report is recorded verbatim as follows:

\# This animal was euthanized early consistent with humane endpoints. However, a necropsy was performed, and tumor cells were analyzed for active numbers and preserved for statistical analysis.

Necropsy Table.

| Group | Observations | Notes |
| --- | --- | --- |
| Control | Bloody ascites, enlarged stomach | Euthanized on day 16 |
| Control | Enlarged stomach | Euthanized on day 16 |
| Control | Enlarged stomach | Euthanized on day 16 |
| Control | Small amount of liquid i.p. | Euthanized on day 16 |
| 8 µg/ml LEV | Slightly bloody ascites, enlarged stomach | Euthanized on day 16 |
| 8 µg/ml LEV | Enlarged cecum | Euthanized on day 16 |
| 8 µg/ml LEV | Small amount of slightly bloodied liquid i.p., enlarged stomach | Euthanized on day 16 |
| 8 µg/ml LEV | Enlarged stomach | Euthanized on day 16 |
| 8 µg/ml LEV | One Hollow Fiber penetrated liver, enlarged stomach | Euthanized on day 16 |
| 16 µg/ml LEV | Bloody ascites, small injuries at the liver | Euthanized on day 13# |
| 16 µg/ml LEV | Highly vascularized vessels | Euthanized on day 16 |
| 16 µg/ml LEV | Cecum filled with gas | Euthanized on day 16 |

Overall, the assay was completed successfully as evidenced by the maintenance of growth of the animals over the 16-day period (Figure 1). Placement of the hollow fiber sections i.p. was observed to cause trauma to some organs; this was the case with the one animal that was euthanized early. There is no evidence that levalbuterol increased the frequency of untoward toxicity since similar findings were found in the control animals that were not exposed to the drug. It is likely that the toxicity was associated with the fact that all animals were carrying the burden of six hollow fiber sections (3 i.p. and 3 s.c.) that contained aggressive lymphoma cultures, which contributed diffusible cytokines, chemokines, enzymes, etc. to the circulation.

## 3. Discussion

Levalbuterol is the levorotary isomer of the racemic drug, albuterol. and has an *R* configuration around a chiral center. It is present in approximately equal quantities of the *S* isomer of albuterol. Most importantly, the *R* isomer possesses all of the broncho dilatory and anti-inflammatory activity of albuterol [1,7]. Also, the *S* isomer is proinflammatory and when present can negate the therapeutic effectiveness and anti-inflammatory activity of the *R* isomer [1,7-11]. The present study only used the pure *R* isomer.

Figures 2 and 3 plotted the descriptive data for the number of active tumor cells in the i.p. control cultures for those animals receiving 8 µg/ml and 16 µg/ml of levalbuterol in the drinking water, respectively. The data illustrates the differences in the proliferation rates for the tumors over the 16 day study period. A treatment concentration of 8 µ/ml did not inhibit tumor proliferation and was not analyzed further. But, 16 µg/ml did affect tumor proliferation. To increase statistical power, the inhibition in proliferation rates for the 3 tumor cultures were normalized to the means of the control cultures before analysis. Two different statistical tests (Figures 4 and 5) were applied and the average percent inhibition was almost identical in both at approximately 55% with strong statistical confidence.

### 3.1. Levalbuterol Potency

Levalbuterol is well absorbed by oral administration. However, from human data, it is known to undergo extensive first pass metabolism by liver and intestinal enzymes that result in only about one-tenth of the absorbed dose reaching the systemic circulation [12- 14]. Specific data for mice are apparently not available. Considering that the mice in this study received levalbuterol ad libitum in the drinking water at 16 µg/ml, and average daily consumption of water for a 20g mouse was 4 ml per day, this would amount to about 3.2 mg/kg body weight intake per day. However, data for areas under the concentration-time curves for the individual ***R*** and ***S*** enantiomers in a cross-over study with 12 young human males gave an estimate of approximately 12% of the administered dose reaching the systemic circulation [12]; this would bring the effective dose to approximately 0.4 mg/kg body weight per day, which is a relatively potent anti-tumor agent.

### 3.2. Supplemental File No.1 Concerning Absence of Classical Glucocorticoid Toxicity with Levalbuterol

A supplemental file taken almost entirely from reference 1 discusses the rationales for the absence of glucocorticoid toxicity with levalbuterol.

## 4. Materials and Methods

Levalbuterol hydrochloride was purchased as the reference standard from the United States Pharmacopeial Convention.

The Hollow Fiber animal assay was conducted by the contract research organization, Reaction Biology Europe GmbH, Engesserstr.4, 79108 Freiburg, Germany.

In this study, female NMRI nude mice (Mus musculus) were used, sourced from Charles River GmbH in Germany. A total of 18 animals, aged 4-5 weeks, were employed. The mice were identified by tattooing and acclimatized for at least 4 days.

The cell cultures involved three cell lines: JEKO-1, MINO, and SU-DHL-1, all derived from human lymphoma. The cells were incubated at 37 °C and 5 % CO2 in RPMI-1640 medium with stable glutamine, 20 % FCS, 100 units penicillin/ml, and 100 µg streptomycin/ml. Routine splitting was performed at a cell density of 1-2 x 10^6 cells/ml for JEKO-1 and MINO, and 0.5-1 x 10^6 cells/ml for SU-DHL-1. Quality control included routine cell line authentication and mycoplasma testing by third parties.

The animals were housed under optimal hygienic conditions in air-conditioned rooms with 10 air changes per hour. The temperature was maintained at 22 ± 2 °C and relative humidity at 45-65 %. The animals were kept in individually ventilated cages (IVC), with a maximum of 4 animals per cage. They were fed M-Zucht diet from ssniff Spezialdiäten GmbH and provided with autoclaved tap water.

On Day -1, tumor cells were loaded into Hollow Fibers and incubated overnight at 37 °C and 5 % CO2. The exact cell numbers for JEKO-1, MINO, and SU-DHL-1 were determined during cell culture.

On Day 0, the mice received subcutaneous analgesia with Meloxicam 1-2 hours before surgery. Hollow Fiber implantation was performed under isoflurane anesthesia, with three fibers implanted subcutaneously and intraperitoneally per mouse.

On Day 16, 14 days after therapy started, the animals were euthanized by cervical dislocation. The fibers were extracted and processed according to the CellTiter-Glo assay protocol. The fibers were homogenized, centrifuged, and incubated with CellTiter-Glo buffer, and luminescence was measured.

The CellTiter-Glo assay is a bioluminescent assay used to determine the number of viable cells in culture. It works by measuring the amount of ATP, a molecule present in all metabolically active cells, using a luciferase reaction. The assay is based on the principle that viable cells contain ATP, and when a reagent containing luciferase and its substrate (luciferin) is added, it triggers a reaction that produces light. The amount of luminescence is directly proportional to the number of viable cells, allowing for accurate quantification. (AI generated.)

GraphPad Prism, Version 10.5.0, was used for all statistical analyses and illustrations.

## 5. Conclusions

Levalbuterol, the pure ***R*** isomer of the racemic drug, albuterol, was found to be a relatively potent inhibitor of proliferation of three human non-Hodgkin lymphomas implanted in mice, in the Hollow Fiber assay model developed at the National Cancer Institute. Two of these lymphomas, JEKO-1 and MINO, are aggressive Mantle Cell lymphomas, the third was SU-DHL-1. Levalbuterol produced an average inhibition of proliferation of 55% for a combination of all three lymphomas. It appears that the mechanism of inhibition is likely apoptosis because levalbuterol is a beta2-adrenergic receptor agonist that shares almost identical gene-expression signatures found in glucocorticoids, which have the capacity to induce apoptosis in malignancies of lymphatic lineage. These findings may have application for possible efficacy in other malignancies such as chronic lymphocytic leukemia, acute lymphoblastic leukemia, multiple myeloma and Hodgkin’s lymphoma.

Levalbuterol is an FDA-approved drug for the treatment of asthma. This study represents an attempt to re-purpose levalbuterol for treatment of neoplastic disorders. The drug is orally bioavailable, has an excellent safety profile and is currently generic, which should facilitate its possible clinical translation.

## Supplementary Materials

Supplemental File No. 1 is attached separately.

## Author Contributions: Conceptualization

funding acquisition; methodology; formal analyses; writing.

## Institutional Review Board Statement

Not applicable.

## Informed Consent Statement

Not applicable.

## Data Availability Statement

All the data are included in the manuscript.

## Acknowledgments

Appreciation is expressed to Jennifer Brown and Gail Anderson of the Department of Pharmacology, New York Medical College, for facilitating administrative and financial procedures required for conduct of the studies.

## Disclaimer/Publisher’s Note

The statements, opinions and data contained in all publications are solely those of the individual author(s) and contributor(s) and not of MDPI and/or the editor(s). MDPI and/or the editor(s) disclaim responsibility for any injury to people or property resulting from any ideas, methods, instructions or products referred to in the content.

---

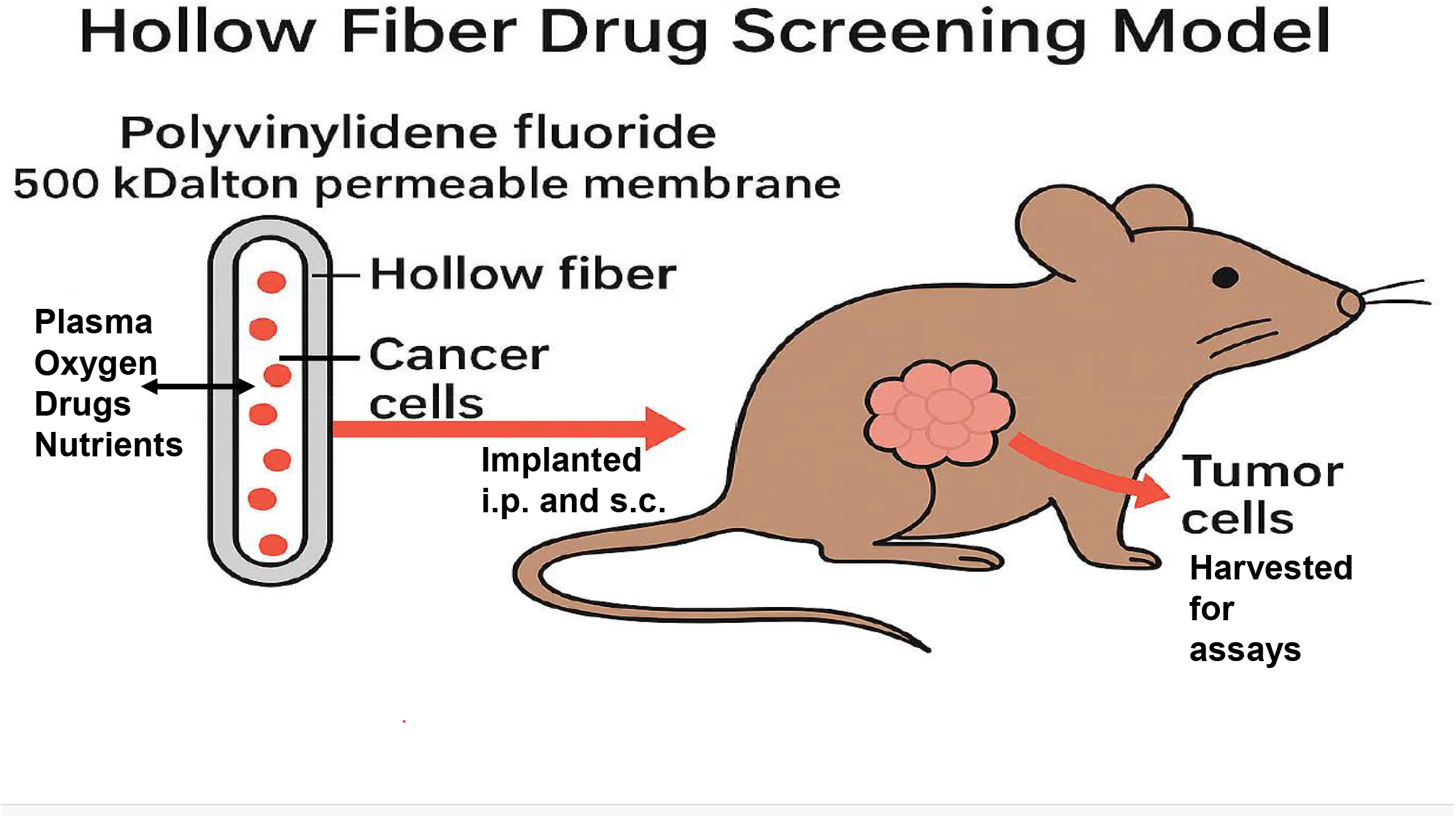

## References

1. Inchiosa, M.A., Jr. Beta(2)-adrenergic suppression of neuroinflammation in treatment of parkinsonism, with relevance for neurodegenerative and neoplastic disorders. Biomedicines 2024, 12, 1720. 10.3390/biomedicines12081720

2. Hollingshead, M.G.; Alley, M.C.; Camalier, R.F.; Abbott, B.J.; Mayo, J.G.; Malspeis, L.; Grever, M.R. In vivo cultivation of tumor cells in hollow fibers. Life Sci. 1995, 57, 131–141. 10.1016/0024-3205(95)00254-4

3. Hall, L.A.; Krauthauser, C.M.; Wexler, R.S.; Hollingshead, M.G.; Slee, A.M.; Kerr, J.S. The hollow fiber assay: Continued characterization with novel approaches. Anticancer Res. 2000, 20, 903–911.

4. Decker, S.; Hollingshead, M.; Bonomi, C.A.; Carter, J.P.; Sausville, E.A. The hollow fibre model in cancer drug screening: The NCI experience. Eur. J. Cancer 2004, 40, 821–826. 10.1016/j.ejca.2003.11.029

5. Spychala, J. Antitumor activity of triazine mimic antibiotics for DNA-binding implications (impressive activity in vitro against a variety of tumor types in the NCI-60 screen): NSC 710607 to fight HCT-116 human colon carcinoma cell lines in vivo using the hollow fiber assay and xenograft mouse models. J. Cancer Res. Clin. Oncol. 2023, 149, 6501–6511. 10.1007/s00432-023-04604-6

6. Montero, M.M.; Domene-Ochoa, S.; Prim, N.; Ferola, E.; López-Causapé, C.; Echeverria, D.; Morisaki, M.F.A.; Vega-Toribio, V.; Sorlí, L.; Luque, S.; et al. Synergistic efficacy of ceftazidime/avibactam and aztreonam against carbapenemase-producing Pseudomonas aeruginosa: Insights from the hollow-fiber infection model. Infect. Dis. (Lond.) 2025, 57, 81–88. 10.1080/23744235.2024.2396882

7. Wang, S.; Liu, F.; Tan, K.S.; Ser, H.L.; Tan, L.T.; Lee, L.H.; Tan, W. Effect of (R)-salbutamol on the switch of phenotype and metabolic pattern in LPS-induced macrophage cells. J. Cell. Mol. Med. 2020, 24, 722–736. 10.1111/jcmm.14780

8. Baramki, D.; Koester, J.; Anderson, A.J.; Borish, L. Modulation of T-cell function by (R)- and (S)-isomers of albuterol: Anti-inflammatory influences of (R)-isomers are negated in the presence of the (S)-isomer. J. Allergy Clin. Immunol. 2002, 109, 449– 454. 10.1067/mai.2002.122159

9. Mazzoni, L.; Naef, R.; Chapman, I.D.; Morley, J. Hyperresponsiveness of the airways following exposure of guinea-pigs to racemic mixtures and distomers of β2-selective sympathomimetics. Pulm. Pharmacol. 1994, 7, 367–376. 10.1006/pulp.1994.1043

10. Chorley, B.N.; Li, Y.; Fang, S.; Park, J.A.; Adler, K.B. (R)-albuterol elicits antiinflammatory effects in human airway epithelial cells via iNOS. Am. J. Respir. Cell Mol. Biol. 2006, 34, 119–127. 10.1165/rcmb.2005-0338OC

11. Ameredes, B.T.; Calhoun, W.J. (R)-albuterol for asthma: Pro [a.k.a. (S)-albuterol for asthma: Con]. Am. J. Respir. Crit. Care Med. 2006, 174, 965–969; discussion 972–964. 10.1164/rccm.2606001

12. Boulton, D.W.; Fawcett, J.P. Pharmacokinetics and pharmacodynamics of single oral doses of albuterol and its enantiomers in humans. Clin. Pharmacol. Ther. 1997, 62, 138–144. 10.1016/s0009-9236(97)90061-8

13. Maier, G.; Rubino, C.; Hsu, R.; Grasela, T.; Baumgartner, R.A. Population pharmacokinetics of (R)-albuterol and (S)-albuterol in pediatric patients aged 4-11 years with asthma. Pulm. Pharmacol. Ther. 2007, 20, 534–542. 10.1016/j.pupt.2006.05.003

14. Hostrup, M.; Jacobson, G.A.; Eibye, K.; Narkowicz, C.K.; Nichols, D.S.; Jessen, S. Beta(2)-adrenergic agonist salbutamol exhibits enantioselective disposition in skeletal muscle of lean young men following oral administration. Drug Test. Anal. 2025, 17, 842–849. 10.1002/dta.3787

