## Supplemental File No. 1 for "Inhibition of Human Mantle Cell Lymphomas by the β_2_-Adrenergic Agonist, Levalbuterol, in the Hollow Fiber Assay in Mice"

**Supplementary Files**

**Supplementary File No. 1**

**Absence of Classical Glucocorticoid Toxicity**

*(Taken from Section 4.1 of the following reference:* Inchiosa, M.A., Jr. Beta2-Adrenergic Suppression of Neuroinflammation in Treatment of Parkinsonism, with Relevance for

Neurodegenerative and Neoplastic Disorders. Biomedicines **2024**, 12, 1720.

https://doi.org/10.3390/ biomedicines12081720)

**Note:** The references have been updated separately for this document.

The marked similarity between gene-expression connectivity for epinephrine (and presumably other β_2_-adrenergic receptor (β2AR) agonists and glucocorticoids, as presented in these investigations, raises the question as to why β2AR agonists have shown absolutely no evidence of the classical adverse effects of glucocorticoids. This is true with chronic treatment of β2AR agonists at pharmacological doses in their primary role as bronchodilators. Adverse side effects of glucocorticoids include hypertension, hyperglycemia, osteoporosis, glaucoma, cataract formation, peptic ulcer, gastrointestinal bleeding, and others [1]. These effects are dose dependent and reduction in dosages leads to loss of therapeutic effectiveness.

Dose limitations represent a major disadvantage in the clinical use of glucocorticoids. In addition to their efficacy as anti-inflammatory/immunomodulatory agents, these drugs have major anti-neoplastic effects in the treatment of several hematopoietic malignancies with lymphatic lineage, including chronic lymphocytic leukemia, acute lymphoblastic leukemia, multiple myeloma, Hodgkin’s lymphoma, and non-Hodgkin’s lymphoma [2,3]. Induction of apoptosis appears to be a primary mechanism in the treatment of these malignancies [4].

There has been considerable effort directed in attempts to separate the desired effects of glucocorticoids from the initiation of adverse events. Agents of this type are non-steroidal chemically and have been variously termed as “dissociated glucocorticoid receptor ligands” [5], “selective glucocorticoid receptor agonists” (SEGRAs) [6], or examples of “biased signaling” [7]. Kleiman and Tuckerman [8] have summarized the results with some of the biological trials with SEGRAs and they have shown limited success.

The fact that epinephrine, the prime example of a β2AR agonist, showed exactly the same gene-expression profile as hydrocortisone in the CLUE analyses but is not associated with any of the adverse effects of glucocorticoids is now known with confidence to be related to the gene-signaling pathways from different receptor ligands. The biological result is related to which gene-signaling pathways are favored [9].

An updated comprehensive review of the glucocorticoid receptor (GR) by Nicolaides et al. [10] discusses the two major classifications of the signaling of gene expression in relation to the potential for adverse events with longer treatment and higher doses versus the pathway for therapeutic predominance. The contrasting pathways are termed “transactivation” and “transrepression.” Transactivation correlates with downstream transcription of protein sequences that mediate the toxic manifestations of glucocorticoid treatment (diabetes mellitus, osteoporosis, hypertension, etc.). Transrepression is primarily associated with therapeutic effects of glucocorticoids, i.e., their anti-inflammatory, immunomodulatory, and certain anti-neoplastic effects. The mechanisms of transactivation and transrepression are complex and are discussed in extensive detail by Lesovaya et al [11]. In view of the potential anti-inflammatory effects of β2AR-related drugs observed in the present study, it would be expected that they mediate via a transrepression effect at the GR.

The chapter by Nicolaides et al. [10] also discusses the evolution of the glucocorticoid receptor, which apparently represents an important step in the survival advantage of a branch that ultimately led to mammalian species. That step, which included specificity of GR for cortisol, took place about 450 million years ago with the first appearance of the GR in ray-finned fish and then to land vertebrates. Of interest to the present study, the evolution of the sympathetic nervous system took place in that same period of geologic time, and it is presumed that a sympathetic nervous system represented a level of functional sophistication that contributed to survival to further evolution [12].

The adrenal steroids and sympathetic nervous systems are critical parts of the integrated response to stress in humans today and this may be the reason that the CLUE platform of the Harvard/MIT Broad Institute database showed essentially the same gene-expression signatures for hydrocortisone and epinephrine, as noted above. In these integrated systems, glucocorticoids have been demonstrated to promptly increase elaboration of cAMP by several mechanisms that would be expected to inhibit the release of pro-inflammatory cytokines and chemokines: ***a***) Hydrocortisone (cortisol) directly activates adenylyl cyclase to elaborate cAMP; ***b***) it increases the density of β2ARs on cell membranes of elements of the innate immune system; ***c***) it facilitates linking of those receptors to adenylyl cyclase to increase cAMP; ***d***) it preserves responsiveness (i.e. reverses tachyphylaxis) that may develop with continuous occupancy of β2ARs [13-18]; ***e***) and finally, there is the added fact that, as part of the stress response, hydrocortisone selectively activates the phenylethanolamine N-methyltransferease (PNMT) enzyme in the adrenal medulla to increase the synthesis of epinephrine [10,19,20]. It is interesting to speculate that the selective beta2-adrenergic-related agents studied in this investigation may extend the clinically therapeutic applications of the glucocorticoid-sympathetic stress response without incurring the adverse effects of higher glucocorticoid doses.

**References**

1.	Schäcke, H.; Döcke, W.D.; Asadullah, K. Mechanisms involved in the side effects of glucocorticoids. *Pharmacol. Ther.* **2002,** *96*, 23–43. https://doi.org/10.1016/S0163-7258(02)00297-8

2.	Isaacs, C.; Wellstein, A.; Riegel, A.T. Hormones and related agents in the therapy of cancer. In *Goodman & Gilman's: The Pharmacological Basis of Therapeutics*, Brunton, L.L., Hilal-Dandan, R., Knollmann, B.C., Eds.; McGraw-Hill Education: New York, NY, 2018; pp. 1237–1247.

3.	Lin, K.T.; Wang, L.H. New dimension of glucocorticoids in cancer treatment. *Steroids* **2016,** *111*, 84–88. https://doi.org/10.1016/j.steroids.2016.02.019

4.	Greenstein, S.; Ghias, K.; Krett, N.L.; Rosen, S.T. Mechanisms of glucocorticoid-mediated apoptosis in hematological malignancies. *Clin. Cancer Res.* **2002,** *8*, 1681–1694.

5.	Schäcke, H.; Rehwinkel, H. Dissociated glucocorticoid receptor ligands. *Curr. Opin. Investig. Drugs* **2004,** *5*, 524–528.

6.	Schäcke, H.; Berger, M.; Rehwinkel, H.; Asadullah, K. Selective glucocorticoid receptor agonists (SEGRAs): Novel ligands with an improved therapeutic index. *Mol. Cell. Endocrinol.* **2007,** *275*, 109–117. https://doi.org/10.1016/j.mce.2007.05.014

7.	Keenan, C.R.; Lew, M.J.; Stewart, A.G. Biased signalling from the glucocorticoid receptor: Renewed opportunity for tailoring glucocorticoid activity. *Biochem. Pharmacol.* **2016,** *112*, 6–12. https://doi.org/10.1016/j.bcp.2016.02.008

8.	Kleiman, A.; Tuckermann, J.P. Glucocorticoid receptor action in beneficial and side effects of steroid therapy: Lessons from conditional knockout mice. *Mol. Cell. Endocrinol.* **2007,** *275*, 98–108. https://doi.org/10.1016/j.mce.2007.05.009

9.	De Bosscher, K.; Haegeman, G.; Elewaut, D. Targeting inflammation using selective glucocorticoid receptor modulators. *Curr. Opin. Pharmacol.* **2010,** *10*, 497–504. https://doi.org/10.1016/j.coph.2010.04.007

10.	Nicolaides, N.C.; Chrousos, G.; Kino, T. Glucocorticoid receptor. In *Endotext*, Feingold, K., Anawalt, B., Blackman, M., Boyce, A., Chrousos, G., Corpas, E., Eds.; MDText.com, Inc.: South Dartmouth, MA, 2000.

11.	Lesovaya, E.A.; Chudakova, D.; Baida, G.; Zhidkova, E.M.; Kirsanov, K.I.; Yakubovskaya, M.G.; Budunova, I.V. The long winding road to the safer glucocorticoid receptor (GR) targeting therapies. *Oncotarget* **2022,** *13*, 408–424. https://doi.org/10.18632/oncotarget.28191

12.	Häming, D.; Simoes-Costa, M.; Uy, B.; Valencia, J.; Sauka-Spengler, T.; Bronner-Fraser, M. Expression of sympathetic nervous system genes in Lamprey suggests their recruitment for specification of a new vertebrate feature. *PLoS One* **2011,** *6*, e26543. https://doi.org/10.1371/journal.pone.0026543

13.	Gross, A.M.; Wolters, P.L.; Dombi, E.; Baldwin, A.; Whitcomb, P.; Fisher, M.J.; Weiss, B.; Kim, A.; Bornhorst, M.; Shah, A.C.; et al. Selumetinib in children with inoperable plexiform neurofibromas. *N. Engl. J. Med.* **2020,** *382*, 1430–1442. https://doi.org/10.1056/NEJMoa1912735

14.	Ciombor, K.K.; Bekaii-Saab, T. Selumetinib for the treatment of cancer. *Expert Opin. Investig. Drugs* **2015,** *24*, 111–123. https://doi.org/10.1517/13543784.2015.982275

15.	Davies, A.O.; Lefkowitz, R.J. Agonist-promoted high affinity state of the beta-adrenergic receptor in human neutrophils: Modulation by corticosteroids. *J. Clin. Endocrinol. Metab.* **1981,** *53*, 703–708. https://doi.org/10.1210/jcem-53-4-703

16.	Parker, C.W.; Huber, M.G.; Baumann, M.L. Alterations in cyclic AMP metabolism in human bronchial asthma. 3. Leukocyte and lymphocyte responses to steroids. *J. Clin. Invest.* **1973,** *52*, 1342–1348. https://doi.org/10.1172/jci107306

17.	Lefkowitz, R.J. Clinical physiology of adrenergic receptor regulation. *Am. J. Physiol.* **1982,** *243*, E43–E47. https://doi.org/10.1152/ajpendo.1982.243.1.E43

18.	Marone, G.; Lichtenstein, L.M.; Plaut, M. Hydrocortisone and human lymphocytes: Increases in cyclic adenosine 3':5'-monophosphate and potentiation of adenylate cyclase-activating agents. *J. Pharmacol. Exp. Ther.* **1980,** *215*, 469–478. https://doi.org/10.1016/S0022-3565(25)32322-0

19.	Adams, M.; Meijer, O.C.; Wang, J.; Bhargava, A.; Pearce, D. Homodimerization of the glucocorticoid receptor is not essential for response element binding: Activation of the phenylethanolamine N-methyltransferase gene by dimerization-defective mutants. *Mol. Endocrinol.* **2003,** *17*, 2583–2592. https://doi.org/10.1210/me.2002-0305

20.	Tai, T.C.; Claycomb, R.; Her, S.; Bloom, A.K.; Wong, D.L. Glucocorticoid responsiveness of the rat phenylethanolamine N-methyltransferase gene. *Mol. Pharmacol.* **2002,** *61*, 1385–1392. https://doi.org/10.1124/mol.61.6.1385
